# Maladaptive response of an endemic California oak to climatic warming is previewed by interannual variation in growth and survival

**DOI:** 10.1101/2024.12.16.628220

**Authors:** Alexander R. B. Goetz, Marissa E. Ochoa, Jessica W. Wright, Victoria L. Sork

## Abstract

Climate change poses a major threat to long-lived tree populations by shifting environmental conditions away from those to which species are adapted. One of the biggest concerns is that individuals that are adapted to current conditions will become maladapted to new conditions due to a reduction in fitness. If tree populations are already maladapted to their current environments, climate change may put them at even greater risk. Valley oak (*Quercus lobata*), a foundational California endemic tree species, has already lost much of its range to anthropogenic activities and has been shown to demonstrate patterns of climatic maladaptation. Given the rapid pace of climate change, predicting the future of a species requires quantifying the extent to which contemporary populations are adapted or maladapted to current climates. Here, we tested the extent and variability of maladaptation in valley oak using a ten-year, range-wide provenance study of 3,371 half-sib juvenile trees from 658 maternal lineages. We compared growth rates and multiplicative fitness functions (height * survival; MFF) across families sourced from sites with differing climates relative to the gardens, as well as against annual phenological records. We found evidence that valley oak trees are most adapted to temperatures beyond the coolest limit of the current species range. Within and across years, trees sourced from hotter localities had higher growth and fitness in the gardens, and the relationship was more pronounced in hotter years. Trees showed persistent maladaptation after ten years, suggesting that the role of phenotypic plasticity in immediate environmental response is superseded by underlying genotypes. Finally, trees with earlier leaf phenology (associated with warmer source climates) consistently showed higher growth rates, further demonstrating a maladaptive pattern. Our results suggest that short-term variation in fitness shows immediate short-term response to climate warming, “previewing” a concerning range of species-level responses to increasing temperature.

## Introduction

Global climate change is jeopardizing the future of ecosystems as climatic conditions shift away from those to which organisms are evolutionarily adapted (Davis and Shaw 2001, Etterson and Shaw 2001, Davis et al. 2005, Jump and Peñuelas 2005). For many species, evolutionary adaptation is expected to be outpaced by rapid climate change within the next hundred years, leading to decreased fitness (maladaptation) relative to historical optima (Carter 1996, Davis et al. 2005, Aitken et al. 2008, Thuiller et al. 2008, Gougherty et al. 2021, Rellstab 2021, Sharma et al. 2022). Evolutionary theory would generally predict that populations within a species are optimally adapted to their current local environment (individuals thus perform best in their home environment, and fitness decreases as they move away from this ideal condition), but not all plants become locally adapted (e.g., Garrido et al. 2012, Etterson et al. 2020, Fréjaville et al. 2020, Gougherty et al. 2021). In this paper, we discuss maladaptation in relative fitness terms (*sensu* Brady et al. 2019), i.e., a population is considered maladapted if its mean fitness value is less than the species’ theoretical maximum fitness. Notably, maladaptation can result from historical adaptational lag (changes in climate outpacing evolutionary adaptation) causing genotypes to be distanced from their optimal climates (Davis et al. 2005, Hendry and Gonzalez 2008, Alberto et al. 2013). If a species is already maladapted, the potential for adaptation outpacing future climate change is likely much lower than in an optimally adapted species, so testing for evidence of maladaptation is necessary to accurately forecast possible future evolutionary trajectories (Rehfeldt et al. 2001, Savolainen et al. 2004, St Clair and Howe 2007). Long-lived tree species are especially likely to exhibit maladaptive patterns due to long generation times constraining genetic recombination (e.g., Frank et al. 2017, Browne et al. 2019, Gougherty et al. 2021, Rellstab 2021, Benomar et al. 2022, Schmeddes et al. 2024). Existing maladaptation, compounded by warming temperatures and habitat fragmentation due to land use change (Kelly et al. 2005), makes it even less likely that trees will be able to outpace climatic changes through either adaptation in place or gene flow (Aitken et al. 2008, Sork et al. 2010). To understand the impact of the extent of adaptation and maladaptation for tree populations, it is valuable to study tree responses across years to interannual fluctuations in climate rather than quantifying average responses across 5 or 10 years with average climate for that or earlier time intervals.

Trees may well experience “decades in which nothing happens and years in which decades happen,” both beneficial and detrimental (Snyder and Ellner 2022). Because the seedling stage is a vulnerable stage of a plant’s life cycle (Collins and Carson 2004, Quero et al. 2008, McLaughlin and Zavaleta 2012), patterns in the first several years of plant growth can have cascading effects that affect the individual’s phenotype later in life (Collins and Carson 2004, Snyder and Ellner 2022). Interannual fitness variation may result from differences in ontogeny (Quero et al. 2008, Barton 2024), which may in turn be either mediated or exacerbated by interannual climate variation (Xu et al. 2022). The timing of bud burst can also vary with temperature across years and thus differentially determine growing season length in genotypes from different lineages (Perry 1971, Lechowicz 1984, Polgar and Primack 2011). As climate change is predicted to increase short-term climatic fluctuations (e.g., more heat waves and droughts; Wang et al. 2017) in addition to increases in average temperatures (Wang et al. 2016, AdaptWest Project 2022), plant response to recent temperature fluctuations can “preview” a range of potential future responses.

In a maladapted species where the climatic optima are already decoupled from local averages, short-term climate fluctuations may subject populations to conditions even further from their optima, potentially leading to pronounced and detectable variation in growth and survival in response. Interannual variation is however rarely a principal focus of longitudinal studies on tree demography as they tend to focus on long-term trends (but see Liao et al. 2022, Schmeddes et al. 2024). Climate change-induced alterations in selective pressures placed on plants will likely not be uniform (Asbjornsen et al. 2021), and aggregate measures may thus conceal impacts of short-term extremes (Dobbertin 2005). Tracking interannual variation in individual growth and survival in common gardens across years thus allows for interpretation of the effects of short-term fluctuations in environmental distance (e.g., the temperature difference between the source and garden may be close to 0 degrees in a cooler year but +3 degrees in a hotter year). Overall, quantification of interannual variation in growth, survival, and phenology among trees allows us to test how plasticity affects short-term fitness outcomes and can in turn empirically demonstrate responses of different genotypes to possible future climate scenarios.

Individual plasticity can also mediate response to rapid climate change where evolutionary adaptation is outpaced (Gunderson et al. 2010, Goessen et al. 2022). In some cases, it can override underlying adaptive genetic variation in determining phenotype (Gimeno et al. 2008, Hamann et al. 2017). If plants are able to adjust to altered climatic conditions, they may show resilience despite genetic maladaptation, but historical plasticity may also contribute to maladaptive patterns by relaxing selective pressures (Hamann et al. 2017). Additionally, individual acclimation and/or differentiation via underlying local adaptation may become more or less apparent at different stages of plant ontogeny (Kueppers et al. 2017, Germino et al. 2019). Acclimation and plasticity thus present a possible “escape route” from maladaptive patterns that, if sufficiently strong, may indicate higher likelihood of successful conservation of valley oak populations. One way to assess the effects of plasticity is to document interannual plant response to interannual climate fluctuations. If phenotypes are highly plastic in traits underlying growth, their growth will be relatively consistent across years with differing climates, but if they are not plastic, we would expect to see declines in growth when short-term climates differ from climates to which the trees are adapted (Alpert and Simms 2002). These data will therefore provide some indication of how plasticity and underlying genetic variation shape tree response to climate.

The overarching goal of this study is to test the hypothesis that individuals are adapted to local climate conditions in an imperiled foundational tree species in California, *Quercus lobata* Née or valley oak, and thus at risk of maladaptation (relative to the species’ fitness maximum) when climate conditions change. Valley oak has already seen major reduction in its range due to land conversion (Kelly et al. 2005), and is predicted to undergo further contraction due to climate change (Kueppers et al. 2005, Sork et al. 2010, McLaughlin and Zavaleta 2012). Using a long-term provenance study of valley oak (Delfino-Mix et al. 2015), Browne et al (2019) found that four-year-old trees growing in two common gardens were adapted to cooler temperatures, suggesting lag-adaptation to historical climates, maladaptation to current climates, and potentially greater maladaptation in the future (Browne et al. 2019). However, those trees were young and possibly still affected by maternal effects (Roach and Wulff 1987). Here, we retest the local adaptation hypothesis and investigate whether evidence of maladaptation persists in ten-year old valley oak trees, given the possibility that the maternal effects will have dissipated, or that the trees have acclimated to local conditions through plasticity (Sultan 1995). Specifically, we will revisit the same individuals in years 6-10 after germination to measure growth and mortality in response to year-to-year climate fluctuations. Additionally, we extrapolate from the empirical models to estimate the hypothetical origin/garden temperature transfer distance that maximizes fitness, which allows us to infer the degree of maladaptation relative to hypothetical historical conditions. Such data will allow us to determine whether maladaptation still drives growth differences across tree lineages and examine how the observed growth trajectories reflect the relative contributions of evolutionary adaptation and individual plasticity to fitness outcomes under climatic fluctuations.

To explore patterns of (mal)adaptation in valley oak in our long-term provenance study, we will test the following hypotheses: (H_1_) Patterns of growth and phenology vary across years due to interannual climate differences; (H_2_) Valley oaks are maladapted today, and the extent of maladaptation varies across the species range; and (H_3_) Evidence of maladaptation decreases over time as trees acclimate to the garden setting. This study will measure two fitness proxies for each garden: Individual relative growth rate (RGR; referred to as “growth rate” in text) in each year and a multiplicative fitness function for each family (height x survival, MFF). RGR allows for standardized comparison of resource acquisitiveness among individuals; here we calculate it on a per-year basis. MFF is an expression of a maternal tree’s level of growth in the common garden that accounts for the percentage of surviving offspring of each maternal tree. We examine the relationships between these metrics and the historical climates experienced by the maternal trees in comparison with the yearly variation in climatic conditions in the common gardens. We then use these relationships to predict theoretical species trait optima in comparison to observed data. To explicitly test fitness differences among families from different environments, we compare RGR and MFF across five cohorts spanning the range of source climates. In addition, we examine how yearly phenology (specifically date of leaf emergence) and source site geography relate to growth and fitness. This study will reassess the earlier evidence that valley oak young trees are better suited for cooler climates than the current climates today and test the prediction that their response to year-to-year fluctuations shows that their response even across years in better in cooler years. We will show that valley oak trees from warmer climates are likely to fare better under climate warming.

## Materials and methods

### Study species

Valley oak (*Quercus lobata* Née) is a long-lived, winter-deciduous tree species endemic to woodlands, savannas, and riparian zones of the California Floristic Province, 0-1700m above sea level (Pavlik 1991). Though its historical range prior to European colonization is estimated to have covered much of the state, up to 95% of the historical range has already been lost to land conversion (Kelly et al. 2005) and only an estimated 3% of its remaining range is within protected areas (Davis et al. 2000). It is also known to be vulnerable to drought and increased temperatures (Kueppers et al. 2005, Tyler et al. 2006, McLaughlin and Zavaleta 2012), but shows evidence of local adaptation to drought (Mead et al. 2019). Furthermore, evidence shows that current valley oak populations are adapted to temperatures from the last glacial maximum rather than the recent past, but the degree of apparent maladaptation differs across genetic lines (Browne et al. 2019). Finally, shifts in local climatic niches of valley oak populations are predicted to outpace gene flow (Sork et al. 2010).

### Common gardens

The findings from this study are based on two common gardens established by J Wright and VL Sork in 2012 as described in Delfino Mix et al (2015). In 2012, over 11,000 open-pollinated acorns were collected from 674 adult trees at 95 localities across the entire species range of valley oak (Figure 1, black dots). Acorns were germinated in 2013, grown for one year in a greenhouse at USDA Institute of Forest Genetics [IFG], Placerville, CA, (Figure 1, blue square) and transferred to lath houses for one year at IFG and the Chico Seed Orchard [CSO], Chico, CA (Figure 1, red square), and then planted into common gardens at IFG and CSO. Trees in the gardens are thus half-sib progeny of the maternal trees. IFG (748m above sea level) has a relatively cool and wet climate, but with hot summer temperatures and below freezing winter temperatures; CSO (69m above sea level) is hotter and dryer than IFG but has higher precipitation and lower winter temperatures than parts of the species range in southern California. In this paper, we refer to these as cooler and warmer gardens, respectively. Trees were planted according to a randomized block design to account for environmental differences between sections of each garden. Environmental differences among blocks (five per garden) are high, especially at the warmer site where two of the blocks are not contiguous with the others. Thus, our analyses account for these differences either statistically or through standardization (see below). Both gardens were irrigated from 2015 to 2019, and the warmer garden was irrigated throughout the entire study period. In 2021, roughly half of the individuals at each garden were thinned according to the original study design to prevent competition. As of 2023, the two gardens include 3,655 surviving juvenile trees from 658 maternal lines.

**Figure 1:**
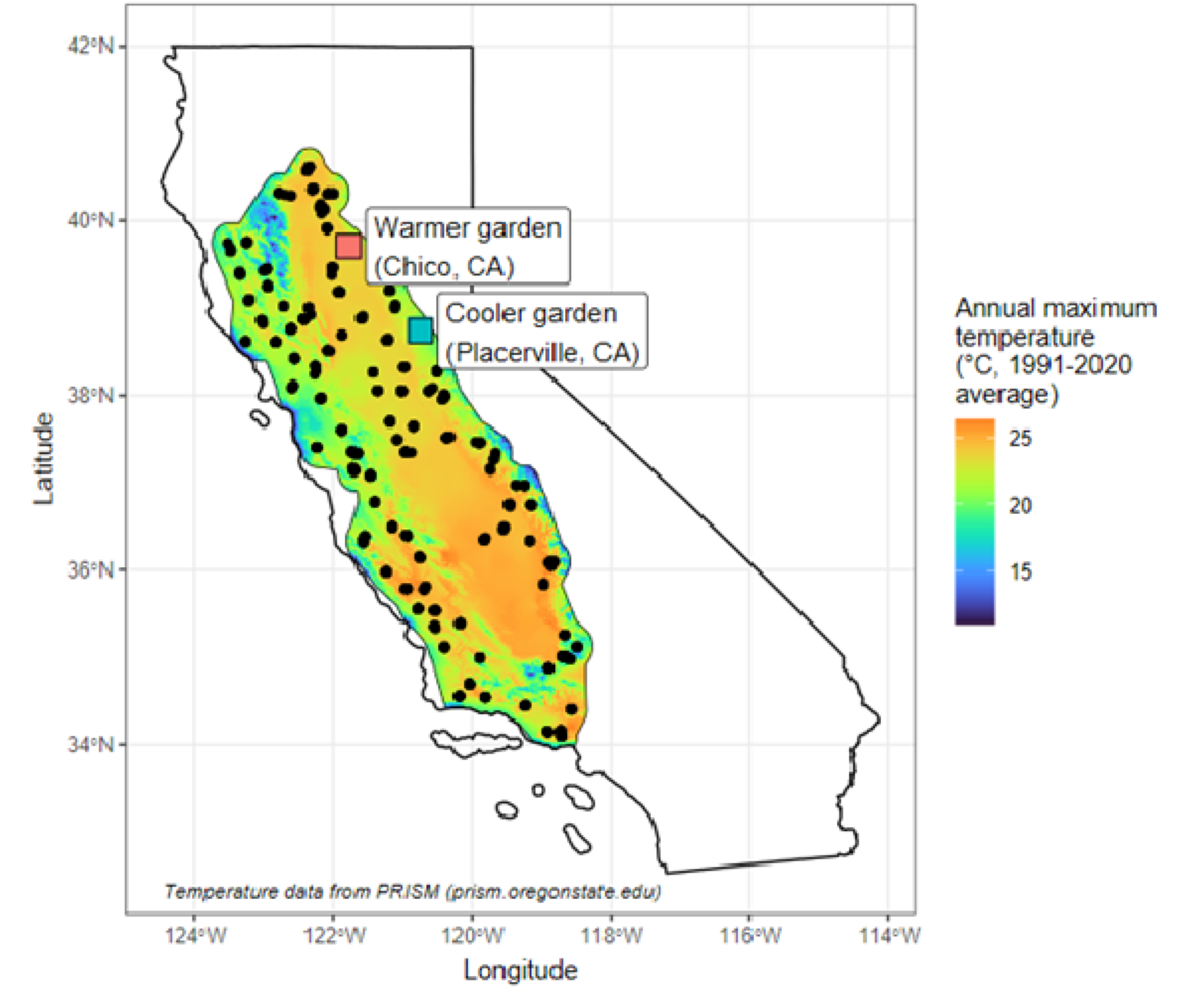
Maternal valley oak tree source locations (black circles) within the current theoretical species range and common garden locations (red square: Chico Seed Orchard, Chico, CA; blue square: Institute of Forest Genetics, Placerville, CA). Background color represents 1991-2020 average of maximum yearly temperature (data from PRISM: prism.oregonstate.edu).

### Survival and growth data

At the end of the growing season in all study years, survival and size were measured (except for part of CSO in 2020 due to COVID-19 restrictions). Survival was assessed visually based on observable presence of living tissue; resprouts of apparently dead individuals were common, so survival data were corrected afterwards. In the early years of the study, size was measured as height of the tallest stem; starting in 2019, trees at least 140 cm tall had diameter at breast height (DBH) measured in addition to height. In 2021, trees removed for thinning had biomass destructively measured. These measures were used to estimate growth. Height was only measured up to a threshold in each year (Table S1), beyond which DBH was used to estimate height, with destructive biomass data used to supplement model training and validation (see below for details). See Table S1 for a detailed list of growth variables measured in each year. In 2018 at the warmer garden, DBH was only measured on a haphazard subset of trees, so we were unable to estimate height for those above the 4-meter threshold; we thus excluded the 2018 warmer-garden data to avoid biases. We also excluded all individuals that showed evidence of herbivore damage (especially a problem in the first year of growth in the gardens), as well as individuals that showed “negative” growth at any point due to either measurement error or dieback. Roughly 200 individuals out of 3,655 were excluded across all years as a result.

### Allometric growth calculations

To allow consistent comparison across different growth measurements, we fit allometric polynomial equations to the relationship between ln(basal diameter) and ln(height) of the trees removed in 2021, resulting in the following formula: *Height* = *^e^*^0.7In(*basal diameter*) + 2.81^ (R^2^ = 0.69). For comparison with other data, we converted DBH to basal diameter using an equation fit based on the relationship between 2018 basal diameter and DBH (basal diameter = 1.42(DBH) + 27.9; R^2^ = 0.72). We then used the basal diameter/height equation to predict height from DBH for trees lacking a true height measurement from 2018-2023. The R^2^ value of the relationship between predicted height and out-of-sample true height was 0.92. All subsequent calculations (growth rate and MFF) were thus based on actual height when present (all individuals 2013-2017; individuals under the cutoff threshold in 2018-2023; individuals haphazardly sampled for complete height in 2023) and height estimated from DBH otherwise. Partially due to inherent patterns of oak tree morphology, the models missed residual differences in the DBH/height relationship across individual trees. Predicted height also saturated at high DBH values. Finally, some trees had multiple dominant stems but only one stem per tree was measured prior to 2023, so predicted height may be underestimated in those cases. These constraints on our measurements of size and growth are unlikely to inflate the significance of our findings, so the trends we observe remain robust. Refer to Table S1 for more details on growth variables across years.

### Relative growth rates

To compare growth across trees from different lineages, we calculated relative growth rates (RGR) for each tree in the common gardens on both an annual and cumulative basis using the following equation:

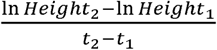

For the annual metric, t_1_ was the year before t_2_. If a tree had missing data for one year, the preceding year was treated as t_1_ (thus, t_2_ – t_1_ was always equal to either 1 or 2 in the interannual comparisons). For the cumulative metric, t_1_ was 2014 (the year before seedlings were outplanted in the gardens) and t_2_ was 2023. Prior to calculating RGR, we excluded any individuals with a recorded decrease in height or estimated height between subsequent years (generally caused by dieback or differences in rounding). We also excluded individuals that had died during the study period or that had been damaged by wildlife. Growth rates were not standardized by block, but differences among blocks were statistically accounted for in all tests by modeling a fixed effect of block nested within garden.

### Multiplicative fitness function by block

To explicitly compare both growth and survival as fitness components of the maternal tree genotypes, we calculated multiplicative fitness functions (MFF) for each family in each garden. Due to known differences in height across planting blocks, particularly at the warmer garden, we calculated individual standardized heights (dividing each tree’s height by the maximum height in its planting block) in each block and averaged across progeny of each maternal tree at each garden. Thus, MFF represents a component of the maternal genotype’s fitness at each garden. MFF was calculated using the following equations (*i* denotes individual progeny in gardens, *j* denotes maternal tree from which all *i* progeny descended).

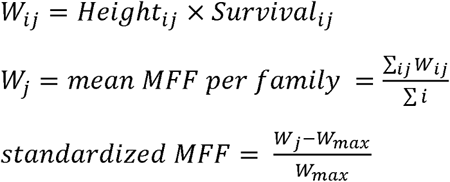

MFF was not calculated for the 2018 or 2020 data at the warmer garden since trees over 4m tall were not measured for height (DBH was only measured on a haphazard subset) and not all blocks could be measured in 2020 due to COVID-19 limitations.

### Temperature transfer distances

To compare the climates of the maternal seed sources to those of the gardens, we calculated transfer distances based on maximum summer temperature for each garden individual in each year, taken as the difference in degrees between maximum summer temperature in each garden each year and maximum summer temperature in the 30-year averages of maternal source climate data. Maximum summer temperatures have been shown to have imposed meaningful selective pressures on valley oak populations in the past (e.g., Kueppers et al. 2005, Sork et al. 2010) and are predicted to increase in the future (Wang et al. 2017). We recognize that precipitation is also an important determinant of valley oak success (e.g., McLaughlin and Zavaleta 2012), but our analyses using precipitation showed the same trends (especially as precipitation and temperature were inversely correlated in the source sites). Moreover, because the gardens were irrigated for during the study period to ensure tree survival for analysis of traits and growth, we are using temperature variables.

Data on maternal tree source site climates from the California Basin Characterization Model (Flint et al. 2013) were used to compare common garden growth outcomes against historical climatic conditions experienced by the maternal source trees. All measurements were 30-year averages of 1950-1981 data gridded at 270m pixel resolution. Monthly climate data for the common garden sites were taken from PRISM (prism.oregonstate.edu). Note that here, unlike Browne et al. (2019), we present the temperature transfer distance in terms of the maternal source climate rather than the planting site (e.g., “trees from warmer climates than the garden” as opposed to “trees planted into cooler environments than the origin”) because it enhanced the clarity of the discussion of our findings. Regardless, the variable’s directionality and interpretation are the same across the two papers.

### Modeling effect of temperature difference-by-year on growth using generalized additive models

We analyzed the effects of fluctuating maternal tree source site climates on growth rate using the generalized additive model framework (GAM; Hastie and Tibshirani 1986, Wood et al. 2015). As we were interested in examining the differences in climate responses across years, the independent variable of interest was the interaction between study year and temperature difference. Maternal tree ID and locality were included as random effects in the models to account for genetic and geographic differences between maternal trees. We also added block within garden as a fixed term to account for microclimatic differences within the gardens. We included initial height of the tree in 2014 as well as the height at the beginning of the interval used to calculate growth, to account for the fact that growth rate inherently decreases with increasing biomass (Rees et al. 2010). All numerical explanatory variables were standardized as Z-scores and modeled using splines with a cubic regression basis. We used Tweedie error distributions for model fitting due to the nonnormal shape of the response variable distribution (Browne et al 2019). The k value (knots used in fitting) was set to 15 for variables that indicated a poor fit with the default value; the default was used for all other variables. We fit GAMs using the bam() function in package ‘mgcv’ (Wood 2011) in R 4.4.0/RStudio 2023.06.1 (RStudio Team 2022, R Core Team 2023). The model was fitted using the following formula:

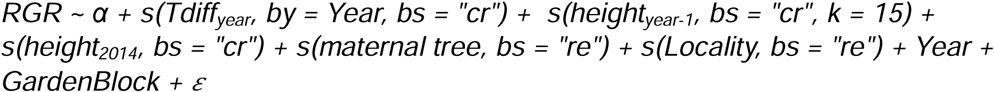

where *α* is the intercept and 𝜀 is the error term; Tdiff_year_ is temperature transfer distance within each year.

### Effects of phenology on fitness

To determine whether phenology influenced growth across years (H_1_), we tested growth rate response to date of first leaf emergence, study year, and temperature transfer distance in each year. The model was fitted using data from both gardens, and included initial height and Block nested within Garden as fixed effects. The model also included source site locality as a random effect to account for regional differences; due to insufficient sample size of phenology data, we were unable to also model differences among families.

Modeling effect of temperature difference on cumulative growth using generalized additive models

To understand the effect of temperature transfer distance on lifetime growth of the trees in the gardens, we also fit generalized additive models based on 2014-2023 growth rate and 2023 MFF (as MFF represents cumulative growth and survival up to the year of measurement). We used the same fitting methods as for the interannual variation models (see above). Models were fitted using the following formulas:

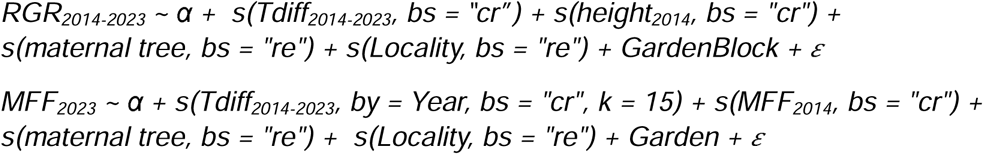

where *α* is the intercept and 𝜀 is the error term; Tdiff_2014-2023_ is temperature transfer distance between the source site average (summer maximum temperature, 1951-1980 mean) and each garden (summer maximum temperature, 2014-2023 mean).

### Extrapolating theoretical fitness maxima: Gaussian process regression

Gaussian process regression is a nonparametric method that allows for extrapolation of models under the assumption that the modeled variable follows a normal distribution (Rasmussen and Williams 2006); Gaussian functions are used in quantitative genetics to mathematically describe the relationship between environment and fitness under local adaptation (Savolainen et al. 2007). To estimate theoretical fitness maxima in relation to the empirical relationship between fitness and temperature transfer distance, we used Gaussian process regression to extrapolate from the GAM fitted values of growth rate and multiplicative fitness. We fit Gaussian process equations to each model (estimating mean and variance based on the shape of the GAM) and predicted values out to a temperature transfer distance of −20°C (the minimum empirical temperature transfer distance was roughly −5°C in the coolest study years). Gaussian process regression was fitted using function gpkm() with a Matern 5/2 kernel in R package GauPro, version 0.2.13 (Erickson 2024).

### Testing for maladaptation: longitudinal cohort study

To explicitly test for climatic maladaptation (e.g., families from warmer source sites than the garden showing higher fitness than families from cooler or similar-temperature source sites), we binned families into five transfer distance cohorts and compared their MFF over time. Transfer distance values were calculated across all observations using the median summer maximum temperature within the study period (cooler garden: 2019, warmer garden: 2023; see Table 1). Cohorts were defined to represent discrete bins of transfer distance values, generally but not always in 2° C increments, to capture local modal transfer distance values as much as possible while also considering relative sample sizes (Figure S1). We then conducted repeated-measures ANOVAs of the cohort*year interaction on MFF with family as a random factor using the aov() function in R package stats. Gardens were analyzed separately since the cohorts were garden-specific. To validate the cohort groupings, we tested several series of similar cohort definitions, including random subsets of individuals. We consistently observed similar outputs, suggesting that the exact cohort definitions do not substantially influence the results.

**Table 1:** Cohorts (A-E) of individuals in two common gardens based on temperature difference ranges (negative temperature difference: trees are from warmer environments than the garden, zero temperature difference: trees are from environments with equal temperature to the garden, positive temperature difference: trees are from cooler environments than the garden). Cohorts are defined differently for each garden (C subscript: cooler garden, W subscript: warmer garden) based on the distribution of temperature differences, resulting in temperature ranges that are not necessarily equal or contiguous between cohorts.

| Garden | Cohort | N | Temperature difference range (°C) |
| --- | --- | --- | --- |
| Cooler (IFG) | A<sub>C</sub> | 64 | (-4.67, -2] |
|  | B<sub>C</sub> | 246 | (-2, 0] |
|  | C<sub>C</sub> | 145 | (0, 2] |
|  | D<sub>C</sub> | 139 | (2, 5] |
|  | E<sub>C</sub> | 59 | (5, 8.9] |
| Warmer (CSO) | A<sub>W</sub> | 31 | (-2.05, 0] |
|  | B<sub>W</sub> | 201 | (0, 2] |
|  | C<sub>W</sub> | 105 | (3, 5] |
|  | D<sub>W</sub> | 109 | (5, 7] |
|  | E<sub>W</sub> | 69 | (7, 9.08] |

### Growth rate comparison across families: norms of reaction

To determine whether differences in growth rates among families changed over time, we conducted norm of reaction analyses that treated each study year as a separate environment. We conducted repeated-measures ANOVAs of the family*year interaction on individuals’ growth rates, controlling for planting block and with individual ID included as an error term, using the aov() function in R package stats.

## Results

### Interannual differences in growth and fitness

To test H_1_, we tested whether year-to-year variation in growth differs due to differences in climate each year. We found that growth rates of individuals differed significantly across years in relation to garden/source temperature difference (Figure 2). For example, in one of the cooler years, 2016, families coming from cooler sites than the gardens show a relative peak in growth rate. However, in most years, the individuals from warmer sites than the common gardens had higher growth rates. Mean growth rate was also variable and tended to be higher in cooler years. The extrapolated relationships show an estimated peak in growth rates close to the temperature transfer distance from the last glacial maximum (−5.2°C) in all but the two coolest years. In most years, predicted growth rate was lower at transfer distance of +4.8°C (the degree of warming predicted by 2100 under the RCP 8.5 “business as usual” scenario) than 0°C transfer distance, but the difference was relatively small (e.g., roughly 5% in 2023). However, in 2021 (the hottest year of the study to date) the predicted growth rate at +4.8°C transfer distance was roughly 19% lower than at 0°C transfer distance. A full model summary is provided in Table S2.

**Figure 2:**
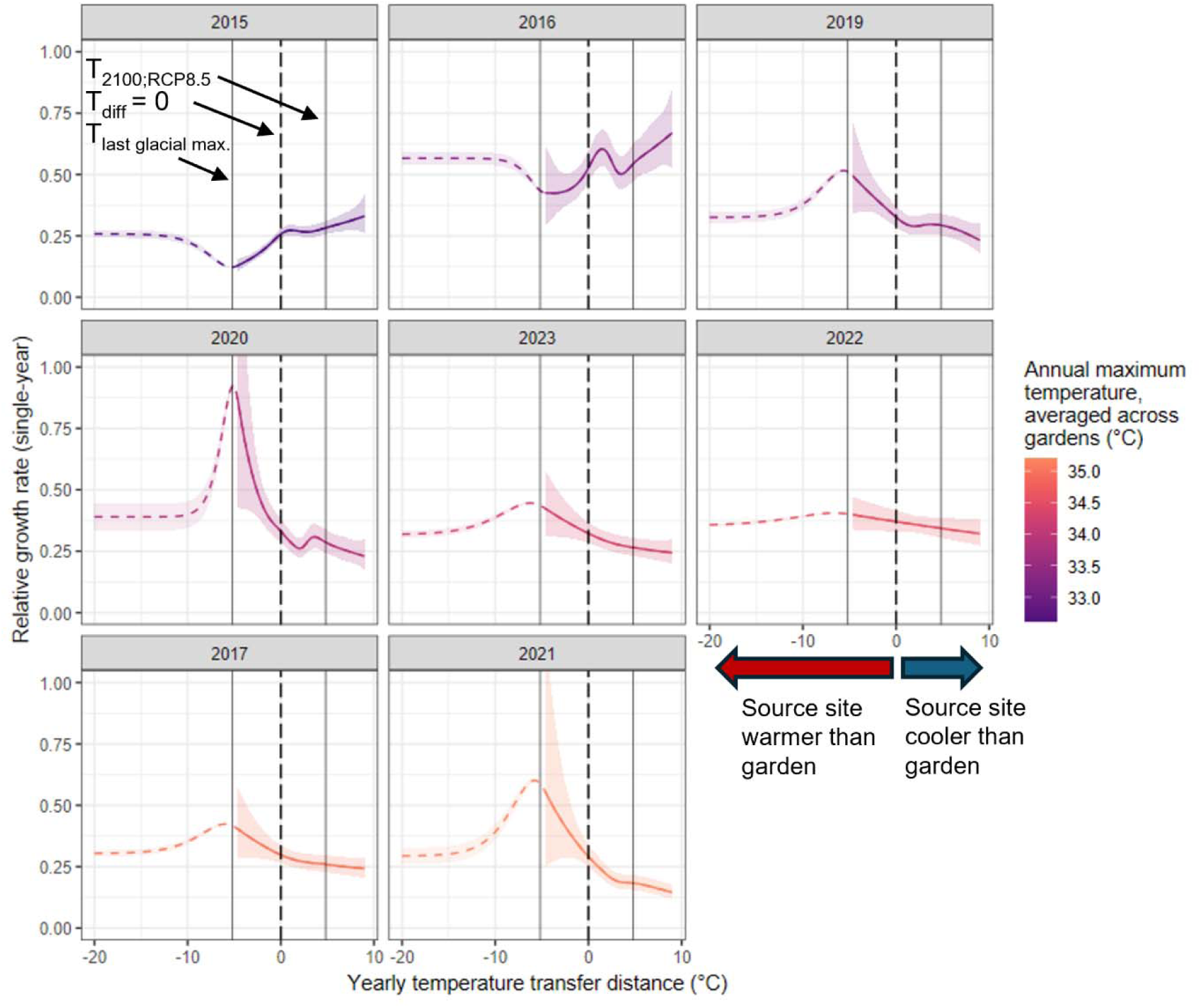
Yearly trends in valley oak growth rate plotted against transfer distance between maternal location and common garden, arranged in order of cooler to warmer temperatures. Solid colored curves represent empirically derived generalized additive model predictions of single-year relative growth rate as a function of yearly temperature transfer distance (based on maximum summer temperature) across the period of tree growth in the common gardens, 2015-2023 (2018 is excluded due to incomplete data). Dashed curves represent extrapolations of the model using Gaussian process regression. Vertical dashed line in each plot denotes where source temperature equals garden temperature; vertical solid lines denote estimated average temperature during the Last Glacial Maximum (21 kya; Wang et al. 2012) and predicted average temperature in year 2100 under a “business as usual” climate scenario (RCP 8.5; AdaptWest Project 2022). The area to the left of the zero line contains individuals from warmer historical climates than the garden, while the area to the right contains individuals from cooler historical climates. Plots are color-coded and ordered by yearly mean temperature across the two gardens.

While we could not directly test the effect of precipitation as plants were irrigated, we observe anecdotally that the shapes of the predicted temperature/growth rate relationships are similar among years with similar mean precipitation, even if temperature was different. For example, mean precipitation across gardens in 2017 and 2019 was 1207mm and 1218mm respectively, but mean temperatures were very different (35.0°C vs. 33.5°C). However, the shapes of the two years’ predicted relationships were more similar than those among the three hottest or three coolest years (Figure 2). Likewise, despite being an “average” year in terms of temperature, 2020 was the driest year of the study (450 mm of precipitation), and had the sharpest predicted fitness decline with increasing distance from the peak (greater penalty for non-optimally adapted genotypes). This would indicate that precipitation may be playing a role in common garden fitness outcomes even despite experimental manipulation of water availability. 2014-2023 annual precipitation at the common gardens is shown in Figure S2.

### Family effects on fitness: Norms of reaction

To directly test the genetic effect of family in relation to interannual variation on growth rate (H_1_), we analyzed the reaction norms of growth rate over time, treating each year’s climate as a separate environment (Figure 3, colored lines). While we could not directly separate the effects of tree ontogeny and climate, increases in temperature between years (Figure 3, black dashed lines) tended to coincide with overall declines in growth rate and vice versa. We also see a large peak in growth rate across all families in 2016, likely due to ontogeny (fast growth at the second-year seedling stage) or acclimation to the site (2016 was the second growing season after out-planting). We found significant effects of year and family, but not year*family interaction, on growth rate at both gardens when controlling for planting block (cooler garden: *F*_family, df = 622_ = 3.0, p < 0.001; *F*_year, df = 8_ = 471.5, p < 0.001; *F*_family*year, df = 4749_ = 1.0, p = 0.39; warmer garden: *F*_family, df = 624_ = 1.7, p < 0.01; *F*_year, df = 8_ = 1286.2, p < 0.001; *F*_family*year, df = 4743_ = 0.79, p = 1). The statistical results indicate that fitness changed among years and families differed in their responses to interannual variation, but their responses did not shift in relation to each other over time (i.e., each family tended to grow either slowly or quickly in all years relative to other families). The results of the norms of reaction are thus consistent with our cohort study, which also found that families tended to separate by fitness in the same way across the study period. Full model summaries are provided in Tables S3 and S4.

**Figure 3:**
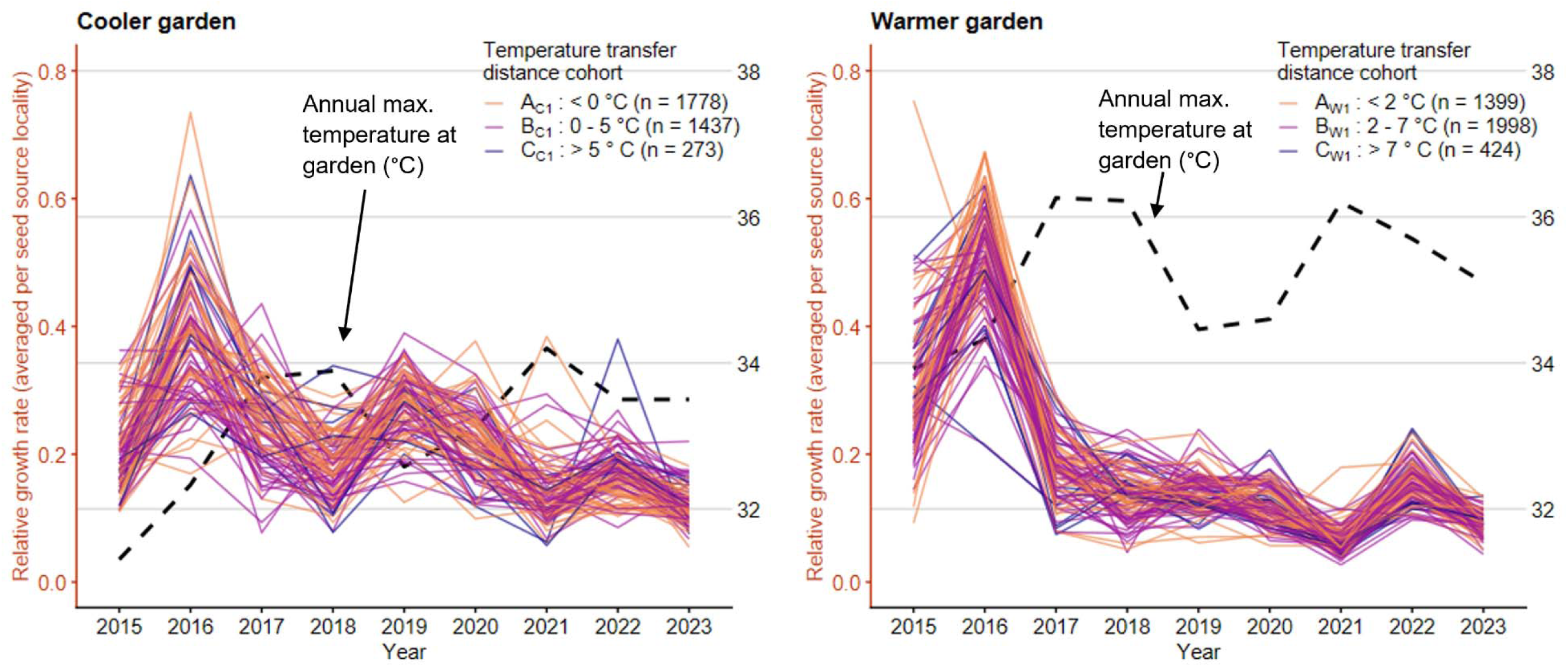
Norm of reaction plots across years for each garden showing average annual relative growth rate of each source site locality (colored lines) and annual garden maximum summer temperatures (thicker dashed black lines). Line colors illustrate cohorts based on temperature transfer distance between garden and source site (indicated in legend; note that these are not the same groupings as in the longitudinal cohort study). 2016 had the highest growth rates on average and relatively low temperatures at both gardens. Likewise, 2017 and 2021 both had increased temperatures and decreased growth rates from previous years. Statistical analysis compares growth rate among families; for easier visualization, each line in this figure represents the average of all families within a geographic locality.

### Cumulative differences in growth and fitness

To determine whether valley oak trees show evidence of maladaptation to temperature after ten years of growth (H_2_), we modeled the effect of temperature transfer distance on cumulative growth rate and MFF. Like the pattern observed above in most study years, we observed faster growth (Figure 4A) and higher multiplicative fitness (Figure 4B) in trees from warmer source sites than the common gardens. The extrapolated relationships from Gaussian regression predict that the theoretical trait optima of growth rate and MFF both occur closer to the transfer distance to the average temperature of the last glacial maximum (−5.2°C) than to a transfer distance of zero (Figure 4A, 4B). Likewise, both trait values are much lower under a predicted future temperature increase of 4.8°C (Figure 4A, 4B). Full model summaries are provided in tables S5 and S6.

**Figure 4:**
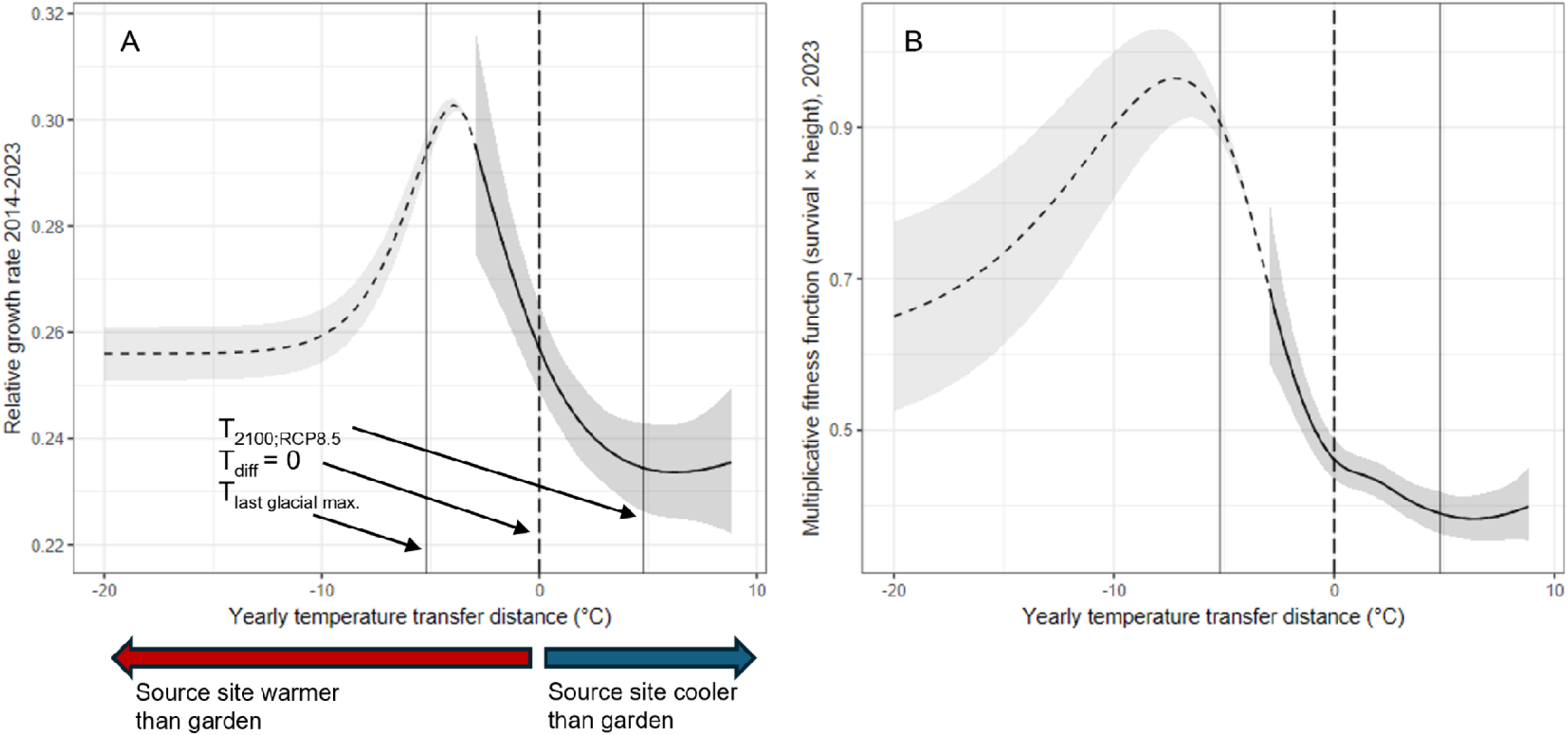
Ten-year cumulative trends in growth rate (**A**) and multiplicative fitness (**B**) plotted against transfer distance between maternal location and common garden, extrapolated to a hypothetical temperature transfer distance of −20°C. Solid black curves denote generalized additive models fit to empirical data, and dashed black curves denote extrapolation of the models using Gaussian process regression. Shaded regions represent 95% confidence intervals. The peak in each extrapolated curve represents the theoretical species-wide trait optimum. Vertical dashed line in each plot denotes where source temperature equals garden temperature; vertical solid lines denote estimated average temperature during the Last Glacial Maximum (21 kya; Wang et al. 2012) and predicted average temperature in year 2100 under a “business as usual” climate scenario (RCP 8.5; AdaptWest Project 2022). The area to the left of the zero line contains individuals from warmer historical climates than the garden, while the area to the right contains individuals from cooler historical climates.

### Cohort study

To test H_2_, we used a cohort approach to test whether individuals with similar source temperatures provided evidence of maladaptation. If populations are maladapted due to lag-adaptation to cooler climates, we expect that families from hotter climates would outperform those from cooler climates. We found significant effects of cohort, year, and cohort*year on MFF of individuals across both gardens (cooler garden: *F*_cohort, df = 4_ = 11.2, *p <* 0.001; *F*_year, df=8_ = 55.7, *p <* 0.001; *F*_cohort*year, df=32_ = 9.5, *p <* 0.001; warmer garden: *F*_cohort, df = 4_ = 2.9, *p* = 0.02; *F*_year, df = 6_ = 76.2, *p <* 0.001; *F*_cohort*year, df = 24_ = 3.6, *p <* 0.001). Tracking MFF of cohorts of individuals (grouped based on their temperature transfer distance in the year with median garden temperature) across the study period also demonstrated 2015-2016 as an inflection point for the emergence of maladaptation; the pattern persisted for the remainder of the study period and the fitness difference between the trees from the hottest and coolest source populations increased over time (Figure 5). In the cooler garden, cohorts B (source site 0-2 degrees C hotter than garden) had the highest fitness in 2015 and a large drop in 2016, after which cohort A (source site 2+ degrees hotter than garden) had consistently higher fitness. Cohorts A, C, and D had relatively similar fitness from 2014-2017, but cohort E consistently had the lowest fitness even in the initial years. Fitness differences among cohorts were more variable across years in the warmer garden than in the cooler garden. Generally, cohorts A and B (cumulatively, source site between 2 degrees hotter and 2 degrees cooler than garden) had the highest fitness across most years and the remaining 3 cohorts had similarly low fitness across years. As in the cooler garden, cohort E (source site 7+ degrees cooler than garden) consistently showed the lowest fitness. In contrast to the cooler garden, the warmer garden showed more consistent fitness responses of all cohorts across years (e.g., all cohorts showed an increase from 2017-2018 and a decrease from 2020-2021). The random effect of maternal lineage explained much more of the variance than the fixed effects (cooler garden: marginal R^2^ = 0.09, conditional R^2^ = 0.75; warmer garden: marginal R^2^ = 0.06; conditional R^2^ = 0.72). Full model outputs are provided in Tables S7 and S8. The two gardens provide two tests of the maladaptation (H_2_), but it is important to note that cohorts were independently defined for each garden and do not represent the same maternal tree individuals across both gardens.

**Figure 5:**
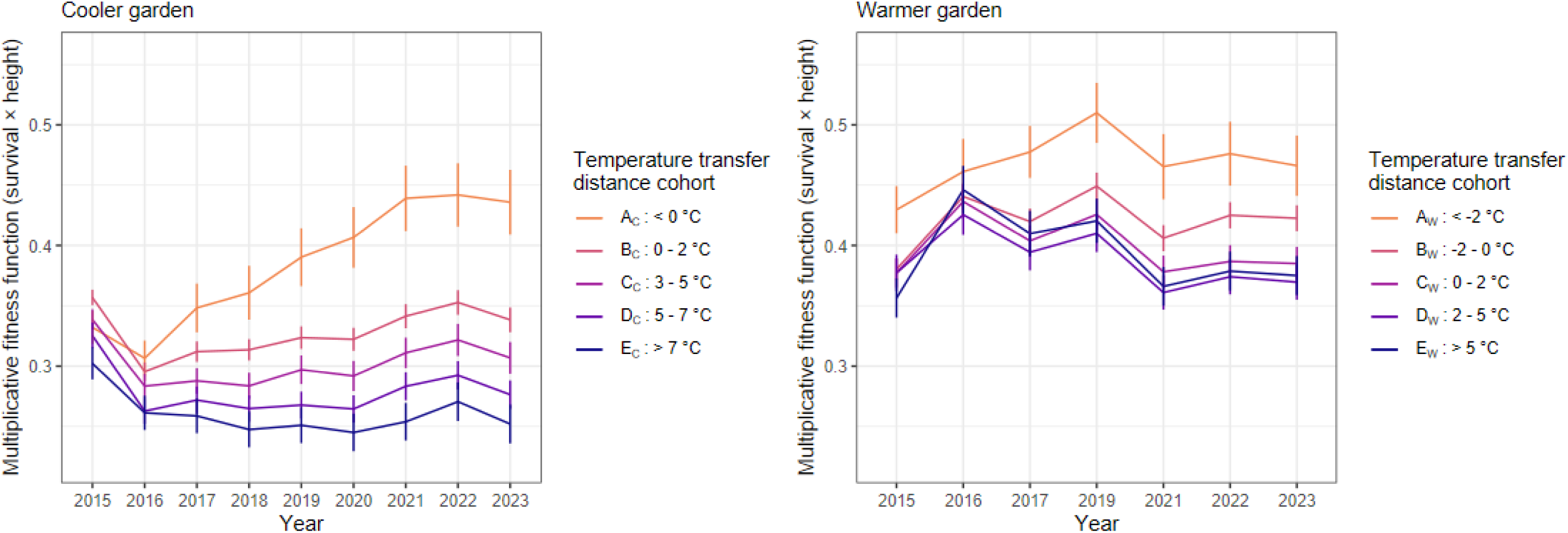
Means and standard errors of multiplicative fitness function of five tree family cohorts at each garden. Differences in MFF by year, cohort, and the year*cohort interaction were all statistically significant (Tables S7 and S8). Each cohort was assigned based on yearly temperature transfer distance between source and garden (e.g., cohort A_W_ consists of plants growing in gardens at least 2°C cooler than their source locations) in the year with median garden temperature (cooler garden: 2019, warmer garden: 2023).

To determine whether phenology predicted fitness outcomes (in particular, whether fitness differences were only due to differences in growing season duration), we tested the relationships between climatic transfer distance, phenology, and growth rate in each year. We observed that families from warmer climates had earlier leaf emergence than those from cooler climates. In addition, relative growth rates were higher in families with earlier leaf emergence across both gardens when controlling for climatic distance between source site and garden (Table 3). In addition, earlier leaf emergence date was associated with higher garden maximum temperatures during early spring when controlling for planting block within garden, year, and maternal seed source (February-April; *p* < 0.001, R^2^ marginal = 0.59, R^2^ conditional = 0.67). These findings indicate that length of growing season contributes to higher growth rates.

**Table 3:**
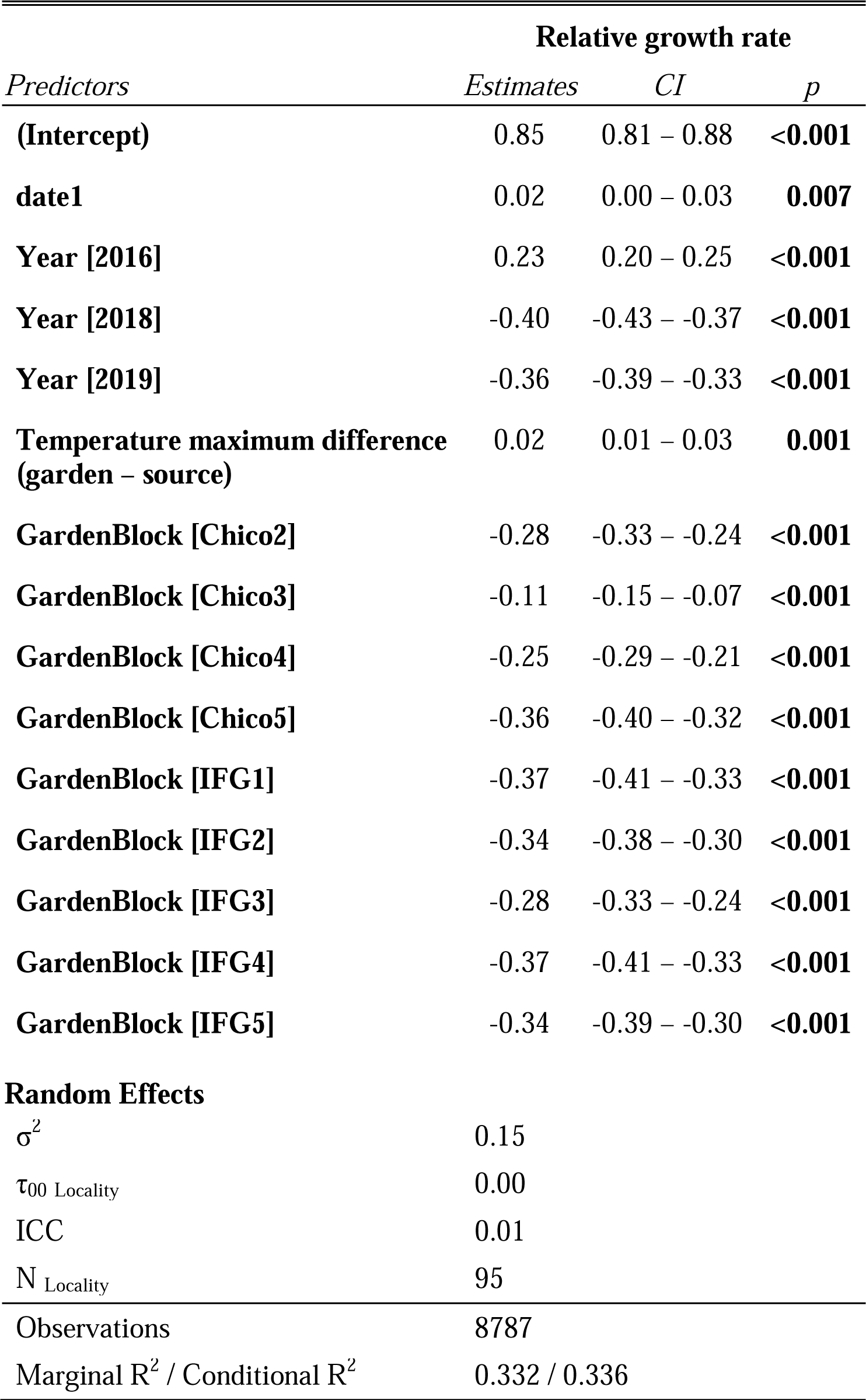
Linear mixed model of valley oak trees’ individual relative growth rates as a function of year, temperature transfer distance, source site locality, and phenology. Planting block within garden was included to account for unintended environmental differences within gardens.

## Discussion

After ten years of growth in common gardens, valley oak trees showed continued evidence of adaptation to climates of 21,000 years ago rather than present-day climates. Moreover, the maladaptive pattern was stronger, and growth rate was lower on average among the entire population, in the hottest study years, suggesting that maladaptation will continue to worsen as average climates warm. Individuals from warmer source sites than the common gardens consistently showed higher fitness than those from cooler source sites, demonstrating that the climate history of maternal lineage was highly predictive of fitness outcomes. Our results point to strong genetic differentiation in response to changing climate, and that trees do not respond with enough phenotypic plasticity to overcome the underlying genotypic maladaptation. The apparent maladaptive trend suggests that valley oaks across their range are likely to suffer reduced growth resulting from increased average and acute temperatures, potentially leading to range contraction and local extirpation in some regions. However, trees from warmer climates may be pre-adapted to future climates, at least in comparison to trees from the rest of the species range.

### Growth rate patterns varied across years (H_1_)

We found strong signatures of interannual variation in metrics of valley oak fitness. In years with higher local temperatures at the gardens, negative relationships between temperature transfer distance and growth rates were more pronounced and growth rates were lower on average. The stronger negative relationship in hotter years supports the maladaptation hypothesis and suggests that populations in the hottest areas of the species range may be close to their climate threshold; in the hottest study year, the temperature at the warmer garden was higher than all maternal seed source historical maxima (i.e., no individuals had temperature transfer distance < 0). In addition, the difference in maladaptive relationship between 2021 and all other years may indicate a threshold of warming tolerance. In most years, the predicted annual relative growth rates between 0°C and +5°C transfer distance were similar, but in 2021 the predicted growth rate was nearly 20% lower among individuals with transfer distance of +5°C. Therefore, we may see not only increased climate mismatch but also increased penalties for climate mismatch as the climate warms beyond trees’ ability to respond to changes.

Our results present evidence that valley oak trees are no longer present in the environments where they should theoretically grow best. Extrapolation of the temperature/growth rate relationships predicted that annual growth rates should peak at a temperature transfer distance of roughly −5°C, which is cooler than any observed transfer distance from a source site within the current valley oak species range. Likewise, given that the warmer garden does not represent a climatic extreme of the valley oak range (Delfino-Mix et al. 2015), the hottest-on-average areas of the species range could become entirely inhospitable to even the most warm-adapted valley oak genotypes if average and acute temperatures rise there. However, climate models predict that cooler inland parts of the species range will experience the most future warming relative to historical conditions (Wang et al. 2016, AdaptWest Project 2022), potentially mediating vulnerability of warm-adapted genotypes and increasing vulnerability of cool-adapted genotypes. While we were unable to directly test the effects of precipitation on growth, we do note an anecdotal indication that the shape of the temperature/growth relationship across years may be associated with precipitation. As water availability has been shown to mediate valley oak response to temperature (Mead et al. 2019), it is likely that both temperature and precipitation will play a large role in determining valley oak fitness under future climates. Nonetheless, our findings demonstrate a clear and immediate danger of climate change not only to the least heat-adapted valley oak populations but to the entire species. As acute maximum temperatures increase over time, the range-wide negative effect of hot years will likely constrain growth of valley oaks in the aggregate (e.g., Sork et al. 2010). We thus find that interannual fluctuations in temperature predict changes in valley oak growth rate over time.

Changes in the relationship between temperature transfer distance and fitness indicated a role of ontogeny in shaping individuals’ responses to climate fluctuations. In the third year of growth in the gardens the direction of the relationship between temperature transfer distance and growth rate reversed from other years. Initially, trees with cooler source climates than the gardens (i.e., positive transfer distance) had the fastest growth, followed by trees with similar source climates to the gardens (i.e., transfer distance around zero). After the second year of growth in the gardens, trees with source climates cooler than the garden consistently grew less per year than those from hotter source climates and the signature of limited local adaptation was no longer present. The fitness difference between hotter-sourced trees and all other trees increased in magnitude for several years and has persisted for the remaining study years. The first two growth years were also the coolest years of the study period, suggesting that short-term fluctuations in local climate have a great degree of influence over short-term growth. However, if these fluctuations were the main driving force behind interannual growth variation we would expect the negative relationship to similarly reverse in subsequent cool years, but in fact the cooler-source trees consistently had lower fitness from 2017 onwards, even in years with similar temperature to 2015 and 2016. These early successes of cool-adapted and locally adapted individuals thus indicate a meaningful ontogenetic distinction within the first few years of growth in which young seedlings show different resource needs than even slightly older saplings (McLaughlin and Zavaleta 2012). The small local maximum growth rate around zero in the second year does suggest some degree of local adaptation; this may be evidence of maternal effects from locally adapted seed sources persisting into the seedling stage and conferring some degree of fitness benefit (Roach and Wulff 1987, Donelson et al. 2018). Indeed, maternal effects were shown to strongly predict early-stage seedling performance in another *Quercus* species (Gimeno et al. 2008). However, any benefit we observed was apparently short-lived (lasting only two years) and has not conferred a detectable lasting advantage to progeny in the gardens.

The interannual fluctuations in growth rate also demonstrate individual phenotypic plasticity across years, but the nonsignificant effect of the family*time interaction indicates that the degree of plasticity is not genetically differentiated and thus not associated with climate of origin (Chambel et al. 2005). Furthermore, the cohort study revealed that individual plasticity was not sufficiently compensatory to overcome the genetic differentiation among families, as those from cooler origins consistently had lower growth despite interannual differences. Directional genetic differentiation overshadowing individual plasticity is consistent with the hypothesis that valley oak trees are lag-adapted as it points to historical local adaptation to varied climates as opposed to species-wide generalization through plasticity (Sultan 1995).

An additional factor that gives trees a benefit in common gardens is the possibility of a longer growing season. As reported previously by Wright et al. (2021), progeny of trees from hotter environments had significantly earlier leaf emergence than trees from cooler environments. We found that earlier leaf emergence was correlated with higher growth rate and relative fitness likely due to a longer growing season. Leaves also emerged earlier on average in years with warmer temperatures during the bud period (February-May), consistent with a previous study on valley oak (Gerst et al. 2017). In turn, we did not observe a tradeoff between a lengthened growing season and risk of frost damage; though late frosts occurred at the cooler garden, we did not observe negative fitness effects of early leaf emergence. This finding suggests a possible future adaptational lag; that is, trees adapted to cooler environments than future conditions may continue to wait longer than necessary to put out leaves and in turn reduce their growth potential. However, the effects of frost damage on other aspects of plant health (e.g., leaf morphology, biomass and nutrient allocation) should be considered as well. We thus find evidence that timing of leaf emergence partially predicts the fitness differences among valley oak trees from differing climates and may be a driver of future shifts toward lower fitness in cool-adapted populations.

As temperatures continue to rise, valley oak trees across their range will be at a considerable adaptational disadvantage relative to their historical optima, but those from cooler source sites are the most negatively affected by high annual temperature increases. Therefore, the “preview” of future climate responses via interannual variation presents a grave scenario for valley oak populations adapted to cooler temperatures than current or predicted future conditions.

### Extent of maladaptation associated with maternal seed source climate (H_2_)

Local climates of the maternal seed source sites were associated with fitness proxies in the gardens, with trees from hotter, dryer, and more precipitation-seasonal climates tending to perform better than trees from cooler, wetter, and more temperature-seasonal climates. Cumulative MFF and growth rate both showed negative relationships with temperature transfer distance, with extrapolated trait optima outside of the current species range. Similarly, MFF varied strongly among cohorts of families from similar source temperature ranges, and the highest MFF was consistently observed in the hottest-adapted cohorts. Instead, variation between maternal tree lineages accounted for most of the explained variance in MFF and significantly predicted between- and among-year variation in growth rate. Previous work on young seedlings in the same study also found that genomic estimated breeding values were more predictive of seedling success than source climate (Browne et al. 2019). Similarly, phenotypic differentiation among families, even those from similar climates, is consistent with other provenance studies (Savolainen et al. 2004, Wilczek et al. 2014, Lind et al. 2024).

The combination of 1) local climate (as opposed to large-scale gradients) as the primary geographic driver of suitability, 2) existing habitat fragmentation (Kelly 2005), and 3) species-wide dispersal and recruitment limitations (Dutech et al. 2005, Tyler et al. 2006) provides a barrier to natural range expansion into cooler areas as climate warms (Savolainen et al. 2004), especially given how severely the families from the coolest parts of the range showed reduced fitness in recent warm years. In turn, our finding of association between fitness and genetics underscores the argument that genotypes in the hottest extents of the species range should be conserved due to their contributions to genetic diversity (Hampe and Petit 2005, McLaughlin and Zavaleta 2012, Richardson and Chaney 2018). In other species, maladaptation is typically driven by factors like genetic drift, gene flow, mutations, and inbreeding (Crespi 2000). Indeed, valley oak gene flow has been shown to be limited relative to historical levels by increased distance between individuals constraining pollen movement (Dutech et al. 2005, Sork et al. 2010). In addition, land use change causing reduction in valley oak range preferentially altered lower elevation sites closer to water as they were more favorable for agricultural and urban use (Kelly 2005), potentially causing genetic drift against genotypes more adapted to these conditions. The entire species is vulnerable to climate change, but cool-adapted populations are particularly poorly suited to current and to future warmer temperatures.

### Limited evidence for acclimation to the common gardens after ten years (H_3_)

After a decade of growth in the common gardens, trees have shown limited acclimation to their planting locations; hot-source individuals continue to show significantly higher fitness than cool-source individuals, but the gap has not widened in the later study years. Single-year relationships between temperature and growth rate were also less steep (i.e., milder fitness penalty associated with distance from the theoretical optimum) in 2022 and 2023 than previous years, but this discrepancy may have been due to climate differences rather than acclimation. While phenotypic plasticity has been observed between trees in the two gardens (MacDonald 2017, Browne et al. 2019, Wright et al. 2021), and now additionally among years, individual plasticity has not been sufficient to overcome the underlying genotypes of the maternal trees. The decade-long persistence of climatic fitness differentiation demonstrates a clear increased risk among individuals across the species range, as genotypes are not locally adapted to their current environments and are thus even less adapted to predicted future climates. The question remains whether the fitness differences among cohorts may eventually decrease, given the long lifespan of valley oak trees. Fitness differences in provenance studies can be dependent on temporal scale--for instance, Germino et al. (2019) only observed local adaptation after 14 years of growth--but other studies (e.g. Kashian and Barnes 2021, Chmura and Modrzyński 2023) found short-term patterns in adaptation to remain consistent across decades. Possible future acclimation will likely also depend on short-term climate, especially as average temperatures continue to rise. The limited but apparent “leveling off” of fitness differences in 2022 and 2023 coincides with both years having maximum temperatures close to the 30-year average; if the gardens experience more 2021-like hot years in the next decade, we might observe more discrepancy in fitness than if temperatures continue to be relatively stable. There is an inherent limitation to our ability to separate the effects of individual plant ontogeny and yearly climatic fluctuations in the gardens, but additional years of monitoring will allow for further decoupling of these factors especially as trees reach maturity. Likewise, our results only take temperature change into account; increased drought due to climate change is also predicted to imperil valley oak populations (e.g., McLaughlin and Zavaleta 2012). Regardless, our findings after a decade of growth in the gardens suggest that early findings of maladaptation (Browne et al. 2019) were not restricted to the early years and were not a result of maternal effects; maladaptive patterns have instead been cemented over time.

## Conclusions

Valley oak as a species shows tremendous vulnerability to climate warming but individuals throughout the species range vary in their vulnerability. Interannual fitness variation among genetic lineages illustrates the sensitivity to tree response of annual climate variation and strong confirmation of range-wide maladaptation. We see anecdotal evidence that precipitation also affects variation in growth, however, the temperature-based analyses are compelling and consistent. Our findings foretell that maladaptation will first negatively impact individuals in the cooler part of the species range, especially where significant warming is predicted to occur, but the entire species will show reduced growth due to increasing temperatures. Furthermore, the relative fitness penalty of maladaptation may increase under warmer climates. Assisted gene flow efforts can aid in conservation (Aitken and Whitlock 2013, Aitken and Bemmels 2016), but choice of genotypes solely based on location or local climate of the source location may not always lead to warm-adapted trees. Instead, strategies involving genome-assisted selection of phenotypes, such as proposed by Browne et al. (2019), may be necessary for sustainability of these populations. We conclude that valley oak populations are already maladapted to current temperatures and thus extremely vulnerable to future climate change.

## Author contributions

VLS and JWW designed the common gardens; VLS, JWW, and ARBG designed research questions; all authors performed research; ARBG analyzed data with input from VLS; ARBG and VLS wrote the paper with input from JWW.

## Acknowledgements

We recognize the native peoples of California as the traditional stewards of the ecosystems in which this research was conducted. We thank Annette Delfino-Mix, Courtney Canning, and Lisa Crane for their assistance in establishing and maintaining the provenance trial. We particularly thank Paul Gugger who led the effort and organized the teams to collect the 11,000 acorns to start the provenance study, as well as many past and present Sork Lab members and volunteers who assisted with data collection. This research was funded by an NSF long-term research grant awarded to VLS and JWW (LTREB-2232794), by funds from the USDA Forest Service Pacific Southwest Research Station, and support to VLS from UCLA.

The findings and conclusions in this publication are those of the authors and should not be construed to represent any official USDA or U.S. Government determination or policy.

Any use of product names is for informational purposes only and does not imply endorsement by the US Government.

