## Supplemental text, figures, and tables for "Maladaptive response of an endemic California oak to climatic warming is previewed by interannual variation in growth and survival"

Supplementary information text

Materials and methods

Maternal tree source climate data

Data on maternal tree source site climates from the California Basin Characterization Model (Flint et al. 2013) were used to compare common garden growth outcomes against historical climatic conditions experienced by the maternal source trees. All measurements were 30-year averages of 1950-1981 data gridded at 270m pixel resolution. Temperature and precipitation data were used to calculate the BioClim variables (worldclim.org), and a principal components analysis was run on 10 climate variables (Maximum summer temperature, minimum winter temperature, maximum, minimum, and average annual temperatures, climatic water deficit, temperature seasonality, precipitation seasonality, precipitation of warmest quarter, precipitation of coldest quarter). The first PC axis (40.8% of variance explained) represented the spectrum between hotter, dryer sites and cooler, wetter sites, while the second PC axis (23.3% of variance explained) represented seasonal differences in temperature and precipitation (higher values indicate temperature-based seasonality, lower values indicate precipitation-based seasonality). For more detail on the climatic variables and PCAs, see Browne et al. 2019.

Effects of geography on growth and fitness

We tested the effects of geography on growth rate and the multiplicative fitness function (MFF). As we were interested in detecting geographic “hotspots” of overall successful families, we measured response of cumulative MFF and growth rate (MFF in 2023; 2014-2023 cumulative growth rate) to latitude, longitude, and the first two principal components describing variation in maternal seed source sites (higher PC1: hotter/dryer, higher PC2: temperature, not precipitation, drives seasonality; see “Maternal tree source climate data” section above). Gardens were analyzed separately. The growth rate models also included the fixed effects of initial height and garden block and the random effect of family. Explanatory variables were scaled as Z-scores. Phenology and geography models were fitted using the lmer() function in R package lme4 (Bates 2010).

Finally, to visualize and interpret the geographic differences in MFF, we used two-dimensional kriging to interpolate MFF values between points within the hypothesized species range. Models were fitted using the gstat() function, and interpolated using the interpolate() function, in R package terra version	1.7-83 (Hijmans et al. 2024).

Results and discussion

Effects of geography on growth and fitness

To test for the effect of geographic variation, including photoperiod, on cumulative MFF and growth rate, we tested the relationship between 2023 MFF/2014-2023 growth rate and longitude/latitude when controlling for the maternal climate PC vectors. We did not find significant relationships between MFF and latitude or longitude (no macroclimate effect), but the PC vectors, representing local climatic variation, significantly predicted MFF. Families from hotter, dryer, and more temperature-seasonal (as opposed to precipitation-seasonal) sites had significantly higher fitness (cooler garden: *p*_PC1_ < 0.001, *p*_PC2_ = 0.01; warmer garden: *p*_PC1_ = 0.02, *p*_PC2_ < 0.001). Geographic variation was high even at the local climate scale, with trees from nearby sites showing varied fitness outcomes. Several maternal trees in the 95^th^ percentile for MFF were sourced from cooler sites than the gardens and many sites hotter than the gardens did not have high performing trees (Figure S3). Geographic effects on cumulative growth rates were slightly different. At the cooler garden, latitude and PC1 both significantly predicted growth rate (*p*_latitude_ = 0.03, *p*_PC1_ < 0.001) as higher in families from hotter sites in the northern extent of the species range. At the warmer garden, only PC2 significantly predicted growth rate (*p*_PC2_ = 0.03), and families from more precipitation-seasonal sites had higher growth rate. Full model summaries are provided in Tables S9-S12.

We did not find strong patterns of latitude or longitude predicting fitness such as reported for forest trees in Canada (e.g., Rehfeldt et al. 1999), nor did individuals clearly separate into distinct geographic units (Liao et al. 2022). Instead, we found high geographic variation in maternal tree fitness. Furthermore, though local climate was a significant predictor of MFF, several trees in the 95^th^ percentile for MFF were from sites up to 2 degrees cooler than the median garden temperature.

Supplementary figures

Figure S1: Distribution of (garden – site) temperature transfer distance values from which cohorts were derived for analysis for the cooler garden (top) and warmer garden (bottom). Bars represent counts of individuals, colors indicate cohorts.

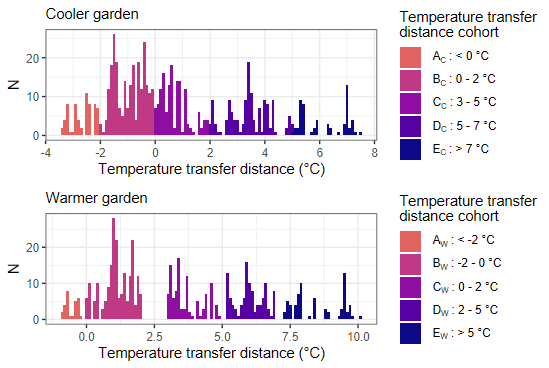

Figure S2: Total annual precipitation 2014-2023 in the warmer garden (red) and cooler garden (blue), with the mean of both gardens shown in the green dashed line. Data from PRISM (prism.oregonstate.edu).

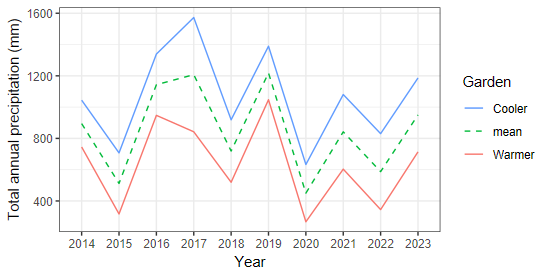

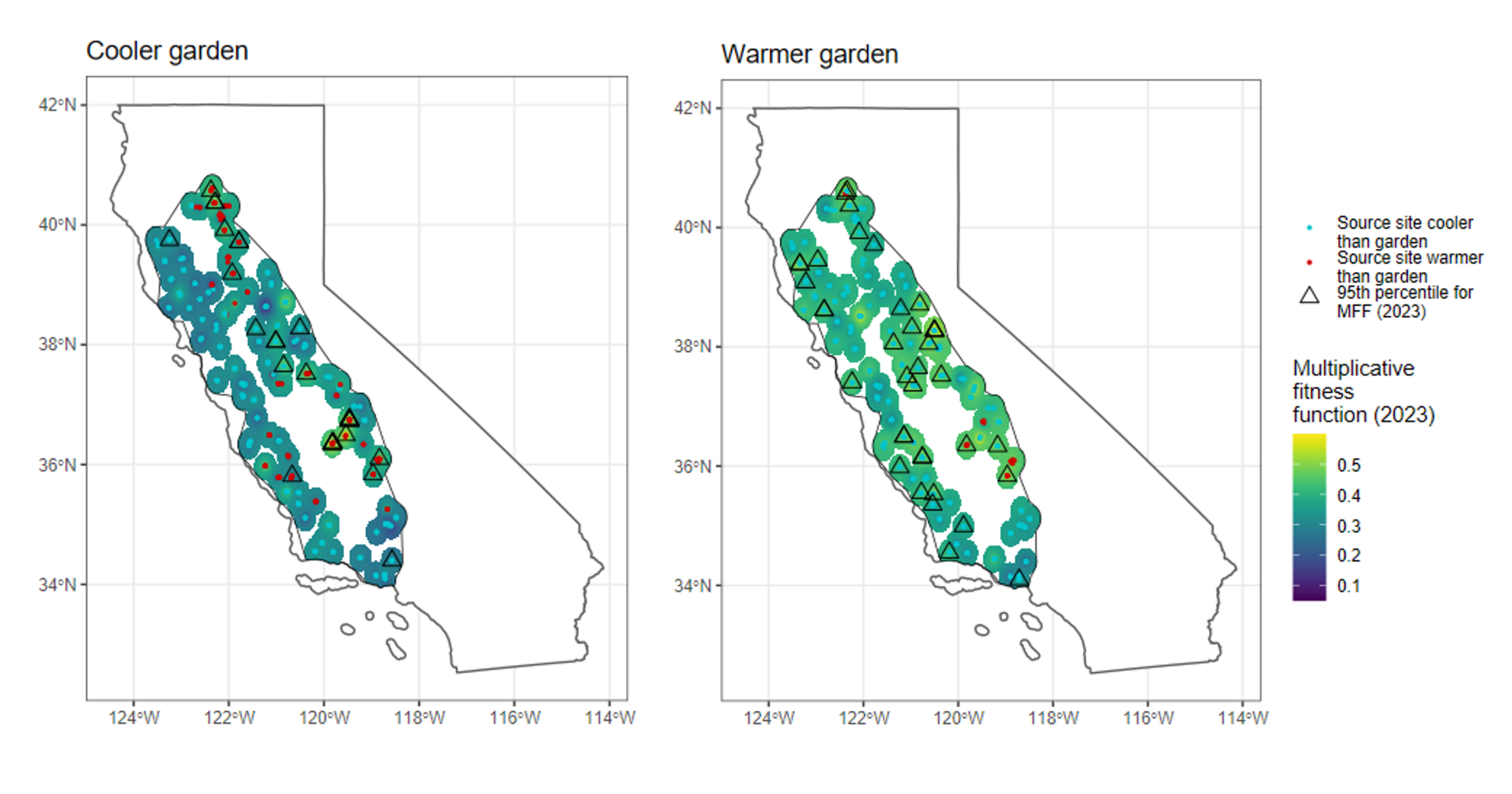

**Figure S3**: Geographic distribution of multiplicative fitness function (MFF) of maternal seed sources measured in the common gardens (left: cooler garden, right: warmer garden) with point locations indicated by blue or red dot surrounded by 2500-meter interpolated buffer zones. Color gradient of buffer zone shows family mean MFF in the common gardens in 2023. The figures also include dots for families from locations with temperatures that are cooler (blue) or hotter (red) than median of maximum summer temperatures in each garden. Black triangles show locations of maternal trees with 95^th^ percentile MFF in each garden as of 2023.

Supplementary tables

Table S1: Summary of growth variables measured on valley oak progeny in the common gardens in each year. Starting in 2018, some individuals were too tall to efficiently measure to their full heights so cutoff thresholds (shown in parentheses) were imposed.

| Year | Growth variables measured |
| --- | --- |
| 2013 | Height, basal diameter |
| 2014 | Height, basal diameter |
| 2015 | Height, basal diameter |
| 2016 | Height |
| 2017 | Height |
| 2018 | Height (<4 m), DBH (haphazard subset), basal diameter (haphazard subset) |
| 2019 | Height (<4 m), DBH (all trees above 150 cm height) |
| 2020 | DBH or basal diameter |
| 2021 | Height (<3 m), DBH (all trees above 150 cm height); basal diameter (removed trees), actual height (removed trees) |
| 2022 | Height (<2 m), DBH (all trees above 150 cm height) |
| 2023 | Height (<2.5 m IFG; <2 m CSO), DBH (all trees above 150 cm height), actual height (haphazard subset) |

Table S2: Full model summary for GAM measuring effects of yearly temperature transfer distance on individual relative growth rates over time.

|  | **Relative growth rate** | | |  |
| --- | --- | --- | --- | --- |
| *Predictors* | *Estimates* | *CI* | | *p* |
| (Intercept) | 0.27 | | 0.26 – 0.28 | **<0.001** |
| Year2016 | 2.08 | | 2.00 – 2.16 | **<0.001** |
| Year2017 | 1.07 | | 1.01 – 1.12 | **0.012** |
| Year2018 | 0.85 | | 0.80 – 0.90 | **<0.001** |
| Year2019 | 1.16 | | 1.10 – 1.23 | **<0.001** |
| Year2020 | 1.15 | | 1.09 – 1.22 | **<0.001** |
| Year2021 | 0.88 | | 0.83 – 0.94 | **<0.001** |
| Year2022 | 1.36 | | 1.29 – 1.44 | **<0.001** |
| Year2023 | 1.12 | | 1.06 – 1.19 | **<0.001** |
| GardenBlock [Chico2] | 0.79 | | 0.76 – 0.83 | **<0.001** |
| GardenBlock [Chico3] | 0.74 | | 0.71 – 0.77 | **<0.001** |
| GardenBlock [Chico4] | 0.69 | | 0.66 – 0.72 | **<0.001** |
| GardenBlock [Chico5] | 0.73 | | 0.70 – 0.76 | **<0.001** |
| GardenBlock [IFG1] | 0.63 | | 0.60 – 0.66 | **<0.001** |
| GardenBlock [IFG2] | 0.61 | | 0.58 – 0.63 | **<0.001** |
| GardenBlock [IFG3] | 0.58 | | 0.55 – 0.61 | **<0.001** |
| GardenBlock [IFG4] | 0.60 | | 0.57 – 0.63 | **<0.001** |
| GardenBlock [IFG5] | 0.55 | | 0.53 – 0.58 | **<0.001** |
| Smooth term (Tdiff_Year_) × Year2015 |  | |  | **<0.001** |
| Smooth term (Tdiff_Year_) × Year2016 |  | |  | **<0.001** |
| Smooth term (Tdiff_Year_) × Year2017 |  | |  | **<0.001** |
| Smooth term (Tdiff_Year_) × Year2018 |  | |  | **<0.001** |
| Smooth term (Tdiff_Year_) × Year2019 |  | |  | **<0.001** |
| Smooth term (Tdiff_Year_) × Year2020 |  | |  | **<0.001** |
| Smooth term (Tdiff_Year_) × Year2021 |  | |  | **<0.001** |
| Smooth term (Tdiff_Year_) × Year2022 |  | |  | **0.003** |
| Smooth term (Tdiff_Year_) × Year2023 |  | |  | **<0.001** |
| Smooth term (Height_Year-1_) |  | |  | **<0.001** |
| Smooth term (Height_2014_) |  | |  | **0.001** |
| Smooth term (Family) |  | |  | **<0.001** |
| Smooth term (Locality) |  | |  | **<0.001** |
| Observations | 23600 | | |  |
| R^2^ | 0.598 | | |  |

Table S3: Full model summary of repeated measures ANOVA on yearly relative growth rate by family (cooler garden).

| Term | df | SS | MS | *F* | *p* |
| --- | --- | --- | --- | --- | --- |
| Error: Individual ID | | | | | |
| Family | 622 | 35.13 | 0.0565 | 3.033 | 5.72e-08 *** |
| Year | 8 | 3.62 | 0.4520 | 24.276 | < 2e-16 *** |
| Block | 4 | 7.72 | 1.9312 | 103.729 | < 2e-16 *** |
| Family*Year | 954 | 46.41 | 0.0486 | 2.613 | 1.20e-06 *** |
| Residuals | 69 | 1.28 | 0.0186 |  |  |
| Error: Within | | | | | |
| Year | 8 | 233.4 | 29.181 | 471.506 | <2e-16 *** |
| Family*Year | 4749 | 296.2 | 0.062 | 1.008 | 0.393 |
| Residuals | 5272 | 326.3 | 0.062 |  |  |

Table S4: Full model summary of repeated measures ANOVA on yearly relative growth rate by family (warmer garden).

| Term | df | SS | MS | *F* | *p* |
| --- | --- | --- | --- | --- | --- |
| Error: Individual ID | | | | | |
| Family | 624 | 52.83 | 0.0847 | 2.115 | 0.000645 *** |
| Year | 8 | 11.22 | 1.4031 | 35.056 | < 2e-16 *** |
| Block | 4 | 14.63 | 3.658 | 91.392 | < 2e-16 *** |
| Family*Year | 1030 | 68.99 | 0.0670 | 1.371 | 0.06980 . |
| Residuals | 51 | 2.04 | 0.040 |  |  |
| Error: Within | | | | | |
| Year | 8 | 1162.6 | 145.33 | 1286.193 | <2e-16 *** |
| Family*Year | 4743 | 422.6 | 0.09 | 0.788 | 1 |
| Residuals | 5493 | 620.6 | 0.11 |  |  |

Table S5: Full model summary for GAM estimating cumulative effect of temperature transfer distance on relative growth rate, 2014-2023.

| **Relative growth rate, 2014-2023** | | |  |
| --- | --- | --- | --- |
| *Predictors* | *Estimates* | *CI* | *p* |
| (Intercept) | 0.26 | 0.25 – 0.26 | **<0.001** |
| SiteBlock [Chico2] | 0.92 | 0.88 – 0.96 | **<0.001** |
| SiteBlock [Chico3] | 0.87 | 0.84 – 0.90 | **<0.001** |
| SiteBlock [Chico4] | 0.84 | 0.81 – 0.87 | **<0.001** |
| SiteBlock [Chico5] | 0.89 | 0.86 – 0.93 | **<0.001** |
| SiteBlock [IFG1] | 0.75 | 0.72 – 0.78 | **<0.001** |
| SiteBlock [IFG2] | 0.72 | 0.69 – 0.75 | **<0.001** |
| SiteBlock [IFG3] | 0.72 | 0.69 – 0.75 | **<0.001** |
| SiteBlock [IFG4] | 0.70 | 0.67 – 0.73 | **<0.001** |
| SiteBlock [IFG5] | 0.64 | 0.62 – 0.67 | **<0.001** |
| Smooth term (Height_2014_) |  |  | **<0.001** |
| Tdiff_2014-2023_ |  |  | **<0.001** |
| Smooth term (Locality) |  |  | **<0.001** |
| Smooth term (Accession) |  |  | 0.956 |
| Observations | 3328 | | |
| R^2^ | 0.608 | | |

Table S6: Full model summary for GAM estimating cumulative effect of temperature transfer distance on multiplicative fitness function, 2014-2023.

|  | **Multiplicative fitness function, 2023** | | |
| --- | --- | --- | --- |
| *Predictors* | *Estimates* | *CI* | *p* |
| (Intercept) | 0.43 | 0.41 – 0.44 | **<0.001** |
| Site [IFG] | 0.74 | 0.71 – 0.78 | **<0.001** |
| Smooth term (Tdiff_2014-2023_) |  |  | **<0.001** |
| Smooth term (MFF_2014_) |  |  | **<0.001** |
| Smooth term (Family) |  |  | 0.577 |
| Smooth term (Locality) |  |  | **<0.001** |
| Observations | 1229 | | |
| R^2^ | 0.223 | | |

Table S7: Full model summary of repeated measures ANOVA on yearly multiplicative fitness function by cohort (cooler garden).

| Term | df | SS | MS | F | p |
| --- | --- | --- | --- | --- | --- |
| Error: Accession | | | | | |
| Cohort | 4 | 6.55 | 1.6372 | 11.202 | **8.62e-09 ***** |
| Year | 2 | 1.08 | 0.5414 | 3.704 | **0.0251 *** |
| Residuals | 643 | 93.98 | 0.1462 |  |  |
| Error: Within | | | | | |
| Year | 8 | 1.830 | 0.22872 | 55.70 | **<2e-16 ***** |
| Cohort*Year | 32 | 1.251 | 0.03909 | 9.52 | **<2e-16 ***** |
| Residuals | 5158 | 21.178 | 0.00411 |  |  |

Table S8: Full model summary of repeated measures ANOVA on yearly multiplicative fitness function by cohort (warmer garden).

| Term | df | SS | MS | F | p |
| --- | --- | --- | --- | --- | --- |
| Error: Accession | | | | | |
| Cohort | 4 | 1.50 | 0.3738 | 2.949 | **0.019866 *** |
| Year | 2 | 2.13 | 1.0627 | 8.384 | **0.000262 ***** |
| Cohort*Year | 2 | 0.09 | 0.0430 | 0.339 | **0.712451** |
| Residuals | 501 | 63.50 | 0.1268 |  |  |
| Error: Within | | | | | |
| Year | 6 | 1.617 | 0.26947 | 76.166 | **< 2e-16 ***** |
| Cohort*Year | 24 | 0.304 | 0.01267 | 3.581 | **9.37e-09 ***** |
| Residuals | 3016 | 10.670 | 0.00354 |  |  |

Table S9: Full model summary of geographic and climatic effects on cumulative family multiplicative fitness function (MFF) in 2023 at the cooler garden.

|  | **MFF** | | |
| --- | --- | --- | --- |
| *Predictors* | *Estimates* | *CI* | *p* |
| (Intercept) | 0.32 | 0.31 – 0.33 | **<0.001** |
| PC1 | 0.05 | 0.03 – 0.06 | **<0.001** |
| PC2 | -0.03 | -0.04 – -0.01 | **0.006** |
| Latitude | -0.00 | -0.03 – 0.03 | 0.869 |
| Longitude | -0.03 | -0.06 – 0.00 | 0.069 |
| Observations | 650 | | |
| R^2^ / R^2^ adjusted | 0.089 / 0.084 | | |

Table S10: Full model summary of geographic and climatic effects on cumulative family multiplicative fitness function (MFF) in 2023 at the warmer garden.

|  | **MFF** | | |
| --- | --- | --- | --- |
| *Predictors* | *Estimates* | *CI* | *p* |
| (Intercept) | 0.40 | 0.39 – 0.41 | **<0.001** |
| PC1 | 0.02 | 0.00 – 0.03 | **0.018** |
| PC2 | -0.03 | -0.04 – -0.01 | **0.003** |
| Latitude | -0.01 | -0.04 – 0.02 | 0.629 |
| Longitude | -0.02 | -0.05 – 0.01 | 0.126 |
| Observations | 629 | | |
| R^2^ / R^2^ adjusted | 0.036 / 0.030 | | |

Table S11: Full model summary of geographic and climatic effects on cumulative relative growth rate across the study period at the cooler garden.

|  | **Cumulative relative growth rate, 2014-2023** | | |
| --- | --- | --- | --- |
| *Predictors* | *Estimates* | *CI* | *p* |
| (Intercept) | 0.29 | 0.28 – 0.30 | **<0.001** |
| PC1 | 0.01 | 0.00 – 0.01 | **<0.001** |
| PC2 | -0.00 | -0.01 – 0.00 | 0.063 |
| Latitude | 0.01 | 0.00 – 0.02 | **0.032** |
| Longitude | 0.00 | -0.00 – 0.01 | 0.372 |
| 2014 height | -0.00 | -0.00 – -0.00 | **<0.001** |
| GardenBlock [IFG2] | -0.01 | -0.02 – 0.00 | 0.227 |
| GardenBlock [IFG3] | 0.00 | -0.01 – 0.01 | 0.710 |
| GardenBlock [IFG4] | -0.01 | -0.02 – -0.00 | **0.006** |
| GardenBlock [IFG5] | -0.03 | -0.04 – -0.02 | **<0.001** |
| **Random Effects** | | | |
| σ^2^ | 0.00 | | |
| τ_00_ _Accession_ | 0.00 | | |
| ICC | 0.08 | | |
| N _Accession_ | 623 | | |
| Observations | 1661 | | |
| Marginal R2 / Conditional R2 | 0.387 / 0.434 | | |

Table S12: Full model summary of geographic and climatic effects on cumulative relative growth rate across the study period at the warmer garden.

|  | **Cumulative relative growth rate, 2014-2023** | | |
| --- | --- | --- | --- |
| *Predictors* | *Estimates* | *CI* | *p* |
| (Intercept) | 0.37 | 0.36 – 0.38 | **<0.001** |
| PC1 | 0.00 | -0.00 – 0.01 | 0.146 |
| PC2 | -0.00 | -0.01 – -0.00 | **0.031** |
| Latitude | -0.00 | -0.01 – 0.01 | 0.993 |
| Longitude | -0.01 | -0.01 – 0.00 | 0.060 |
| 2014 height | -0.00 | -0.00 – -0.00 | **<0.001** |
| GardenBlock [Chico2] | -0.03 | -0.03 – -0.02 | **<0.001** |
| GardenBlock [Chico3] | -0.03 | -0.04 – -0.02 | **<0.001** |
| GardenBlock [Chico4] | -0.05 | -0.05 – -0.04 | **<0.001** |
| GardenBlock [Chico5] | -0.04 | -0.05 – -0.03 | **<0.001** |
| **Random Effects** | | | |
| σ^2^ | 0.00 | | |
| τ_00_ _Accession_ | 0.00 | | |
| ICC | 0.01 | | |
| N _Accession_ | 625 | | |
| Observations | 1718 | | |
| Marginal R^2^ / Conditional R^2^ | 0.549 / 0.553 | | |
